# Reconciling the importance of minerals for propagation of antibiotic resistance genes in the environment

**DOI:** 10.1101/2024.03.04.583397

**Authors:** Saghar Hendiani, Carlota Carbajo Moral, Mads Frederik Hansen, Oluwatoosin Bunmi Adebayo Agbaje, Pablo Nicolas Arellano Caicedo, Taru Verma, Ines Mandić Mulec, Mette Burmølle, Karina Krarup Sand

## Abstract

The role of mineral surfaces in environmental processes, particularly their influence on DNA preservation, biofilm formation, and genetic transfer, has garnered attention due to its implications for the spread of antibiotic resistance genes (ARg). Despite the recognized significance of mineral-mediated DNA transfer, this mechanism remains poorly understood. Here we investigate the intricate interplay between soil minerals, bacteria, and DNA, to better understand the mechanisms driving ARg propagation in natural environments. We here study the uptake of mineral adsorbed DNA into the natural competent bacteria *b. subtilis* and further explore the influence of minerals on the viability and subsequent biofilm formation of both *b. subtilis* and *A. baylyi.* We further adsorbed DNA to mineral surfaces and allowed biofilm formation while monitoring the propagation of the ARg through out the biofilms. All the results are set in context of mineral surface properties such as surface charge, charge densities and surface area.

Our results showed that the surface properties of the mineral surfaces are highly influencing the transformation efficiencies, viability and biofilm formation where in particular a high number of positive charged surface sites enhance biofilm formation and viability and inhibit transformation. The influence of the mineral surfaces diminishes as the biofilm develops and propagation of mineral adsorbed ARg are seen widely across the mineral surfaces. Our results have implication for mitigations strategies and reconcile mineral surfaces as hot spots for the propagation of antibiotic resistance-which indeed can be driven by transformation in the absence of bacteria carrying the traits. In principle all it takes is one successful transfer event from a mineral adsorbed ARg.

## INTRODUCTION

It is well established that mineral surfaces play a role for DNA preservation in the environment, biofilm formation and development, and in DNA transfer from mineral surfaces. Subsequent propagation of mineral adsorbed DNA has received little to no attention despite its implications have been raised [1-3]. Here we aim to reconcile that the minerals have a strong influence on biological processes in our ecosystems and showcase implications for the environmental propagation of Arg.

The spread of antibiotic resistance genes (ARg) is a worldwide health risk [4] and is no longer only a clinical issue. Vast reservoirs of ARg are found in natural environments [5-7] such as soils, sediments, and oceans. The release and dissemination of ARg to the environment is primarily caused by extended use of antibiotics in farming, where the genes dissipate from the manure [8]. Once in the environment, the ARg are surprisingly rapidly propagated. It is well known that the ARg are distributed to neighbour bacteria through processes of both cell division or through horizontal gene transfer (HGT) where one species acquires resistance from another [9, 10]. Most HGT responsible for the spread of ARg is assumed to be through direct microbe-microbe contact. However, recent work on mineral facilitated HGT indicate that the non-contact transfer mechanism is grossly underestimated. In the HGT mechanism called “Transformation”, free ARg in suspension or ARg adsorbed to mineral surfaces can be picked up and incorporated into distantly related bacteria. Considering that free DNA only persists for a few weeks in sea- and freshwater environments before being degraded,[11-13] any transformation from free DNA can rightly be assumed to be local. However, recent findings show that if DNA is associated with sediments, it can survive for up to 2 Ma[14]. In contrast to the short longevity in seawater minerals offer a potent pathway for preserving and propagating ARg across time scales and across environmental systems. Given that many bacteria anchor themselves to mineral surfaces and that only a few sedimentary systems are void of bacteria, we argue that mineral surfaces are a much-overlooked pathway for shuttling ARg to distant organisms and ecosystems. Once bacteria are in contact with the mineral, they can uptake the adsorbed DNA and the mineral can further act as a platform for spread of ARg to multiple organisms through successions of cell division and HGT. The ability of mineral surfaces to act as hotspots for genetic exchange is much understudied and more knowledge of drivers in these processes could help improve mitigation strategies to prevent ARg propagation in our environments.

Transformation of ARg from minerals is little explored, but results suggest that the interactions between bacteria and minerals [15] as well as interactions between DNA and mineral [16] may play a role for uptake frequency. Huang et al. investigated how kaolinite, montmorillonite, and goethite affect *B. subtilis* transformation. Kaolinite and montmorillonite decreased transformability by adsorbing a cell competence signaling molecule. In contrast, goethite in higher concentrations caused significant membrane damage, leading to a notable increase in bacterial transformation. [16] Taru et al recently showed that fragmented DNA could be taken up from common mineral surfaces of clays (kaolinite and mica) iron oxides (hematite and goethite) carbonates (calcite) and a non-clay silicate (quartz)[3]. As also observed in the studies of ARg plasmids, the rate of transformation of fragmented DNA varied across the minerals. Taru et al. showed that the DNA-mineral association caused the observed differences in uptake efficiency where a strong DNA-mineral association led to a decrease in uptake frequency [3].

Minerals have distinct surface properties as controlled by their crystalline structure and composition, and their surface charges most commonly vary with pH. Double stranded DNA molecules mainly interact with minerals through their phosphate backbone, with a small contribution from nucleobases [17, 18]. The pPhosphate group on the backbone is negatively charged in most environmental conditions and can bind directly to positively charged mineral surfaces (such as oxides and clay edge-sites). In contrast, binding of DNA via the phosphate backbone groups will require cationic bridges in order for the DNA to bond to negatively charged mineral surfaces (such as quartz and clay basal planes). The DNA adsorption capacities of minerals vary widely but can generally be described by interfacial geochemical principles. Consequently, the mineral surface characteristics will in an interplay with the environmental conditions determine the DNA-mineral affinity and stability [19, 20], [21-25], [24, 26-32]. So far, most studies of DNA-mineral interactions focus on bulk level interactions, but also spectroscopic [23, 26, 33, 34] and QCM-D approaches have been made [19, 20, 33, 35, 36]. Recent nano-level investigations have showed that the strength of the DNA mineral association depends on mineral surface topography and the active site density for binding with the molecule in questions [18]. Verma et al showed that the binding affinity between the DNA and the mineral surface and the mobility of the adsorbed molecule scaled with transformation efficiency which highlights that nano-level insight into DNA-mineral binding is an important parameter to consider obtaining mechanistic insight about mineral facilitated transformation [3].

Atomic force microscopy (AFM), where a nm sized probe rasters across a surface, has establish that the negative surface charge of a mica mineral is not able to scan the full extent of an adsorbed DNA molecule. Instead, a high charge dense cation such as Mg or Ni needs to be applied to provide a bond strong allow for imaging. In the presence of Mg, the adsorbed plasmid can be viewed in air where it displays circular strands with a few twists of the helix. If imaged in liquid the DNA molecule is only showed as a few moving specs on the scanned images as the bond between the molecule and the surface (incl. Mg ions) is not sufficiently strong to retain the DNA to allow imaging. AFM has also been used to study interactions between DNA and the (104) face of calcite, (001) face of hematite and (001) face of chlorite. Chlorite and calcite both display topography on the surface and the AFM studies highlight how the associated charge and adsorption site densities change with the local topography. I.e. Verma et al showed that short DNA fragments adsorbed to calcite step edges show no mobility while imaging in liquids but that strands adsorbed to the flat terraces showed mobility [3]. For plasmids, Freeman et al, showed that the adsorption is dominated by the charge dense step edges where the DNA molecules show little mobility [18].

Besides being instrumental for DNA-mineral interactions, the mineral surface properties have also been shown to play a role for the ability of bacteria to form biofilms [37-39]. In the initial steps in biofilm formation bacteria-mineral interactions play a pivotal role influencing bacterial attachment and biofilm formation. Bacterial cells possess charged components on their surfaces, affecting their affinity for different surfaces. Electrostatic forces between the cell envelopes and the mineral surface determine initial attachment, guiding the bacteria to adhere or repel [40]. Opposite charges often promote adherence, facilitating biofilm initiation and growth. i.e. Stenström has shown that surface charge and hydrophobicity of minerals play crucial roles in modulating bacterial adhesion [41, 42]. Additionally, surface charge impacts biofilm architecture, influencing its stability and resilience against environmental stressors [42, 43].

Interactions between bacteria and mineral surfaces are also shown to affect bacterial viability, metabolic activity, competency, and the structure of biofilm aggregates [16, 44, 45]. In particular, metabolic activity and viability is affected by the electrostatic interactions and adhesion forces between bacterial cells and mineral surfaces [46]. The mineral goethite causes cell death caused by a strong interaction between cells and the goethite surface [47, 48]. and Huang et al showed that the interaction with goethite also caused a preference for upregulating motility [47]. Shi et al showed that adsorption of the plasmid by minerals would change plasmid morphology and cause changes in bacterial response in DNA uptake [49]. Also, some minerals can influence the bacterial cells by enhancing membrane permeability and DNA uptake through production of reactive oxygen species (ROS) [50] [49]. A general consequence of membrane damage is increased stress response which can lead to enhanced biofilm formation. Mineral surfaces can also indirectly modulate gene expression by adsorbing signalling molecules and thereby e.g. decrease bacterial competency reducing transformation efficiency [51]. Furthermore, minerals are shown to significantly impact the biofilm structure by influencing production of matrix extracellular polymeric substances (EPS)and matrix structure and composition, i.e. mineral interactions with biofilms can lead to alterations in microbial communities, their structure, and functions. Biofilms formed on different minerals undergo alterations in architecture, which can change the spatial structure of bacterial association in biofilm matrix [43]. This restructuring can influence the distribution of nutrients, signaling molecules, and genetic material exchange within the biofilm, ultimately impacting microbial activities and biogeochemical processes [52]. Further adsorption to mineral surfaces has been shown to change biofilm protein function [53], directly affecting various metabolic reactions, including signal transduction [54], cellular adhesion [55], stress responses [56] and EPS formation [43].

We use two common soil bacteria naturally capable to uptake foreign DNA (Naturally competent) as model organisms: the Gram-positive *Bacillus subtilis* and the Gram-negative *Acinetobacter baylyi* and a broad range of soil minerals (iron oxides, clay minerals, carbonates, silicates, and quartz). *B. subtilis* is spore-forming, flagellated and motile and is commonly used in studies of transformation, conjugation, and biofilm formation. The *B. subtilis* adapts to various ecological niches and is found in diverse habitats from soil, water, and air to the gastrointestinal tract. *A. baylyi* is widely applied as a model organism because of its metabolic versatility and genetic plasticity and unlike *B. subtilis*, *A. baylyi* does not form spores but demonstrates robust resistance to environmental stress. It possesses pili as appendage which are used to facilitate surface attachment. *A. baylyi* is commonly found in diverse habitats, including soil and water. The biofilm matrix of both bacterial strains includes polysaccharides, proteins, and extracellular DNA. *A. baylyi* biofilms typically exhibit thick matrix, aiding in attachment and protection, while *B. subtilis* biofilms are characterized by their extensive matrix restructuring to be used mostly for nutrient acquisition [57]. One notable difference in the biofilm matrix of *A. baylyi* is its high content of exopolysaccharides, which contributes to structural integrity and water retention [58]. In contrast, *B. subtilis* biofilms are characterized by a matrix rich in amyloid-like fibers called TasA, aiding in surface adherence and cohesion [59]. *A. baylyi* biofilms often display higher resistance to environmental stresses due to specific genetic adaptations, whereas *B. subtilis* biofilms demonstrate greater architectural complexity for enhanced tolerance. Moreover, *A. baylyi* biofilms display increased adhesion to surfaces facilitated by pili, whereas *B. subtilis* relies more on exopolysaccharides for attachment. The regulatory networks governing biofilm formation differ between the two species, with distinct signaling pathways and transcriptional regulators identified [60, 61].

We used 2 types of nano particulate iron oxides: goethite and hematite, 2 types of clay minerals: muscovite and kaolinite. Muscovite (mica) is merely representing a clay basal plane where kaolinite is a common clay mineral composed of alternating tetrahedral (as mica) and octahedral oxide layers (Si_2_O_5_ and (Al_2_(OH)_4_) respectively). The aluminum oxide/hydroxide layers in kaolinite are also called gibbsite layers. We used calcite (CaCO_3_) to represent carbonates and quartz to represent non-clay silicates. We used bulk derived parameters of DNA adsorption capacity, surface area and site densities as well as nano-level derived information of the mobility of adsorbed DNA as key parameters in our analyses of the effect of the mineral surfaces for transformation, viability, biofilm formation and subsequent ARg propagation in the biofilms. Antibiotic selective agar plates to quantify the rate of transformation. Atomic force microscopy (AFM) was applied to image DNA-mineral interactions. Viability of bacteria exposed to mineral were measured using colony forming units (CFU/ml), combined with LIVE/DEAD staining under fluorescent microscopy for membrane integrity. We measured the metabolic activity of biofilm buildup using TTC and used scanning electron microscopy (SEM to characterize biofilm morphology.

## RESULTS

### Uptake by transformation of mineral adsorbed DNA

We observed marked differences in transformation efficiency of *B. subtilis* among the six minerals (Fig. 1). The negatively charged mineral, quartz displayed the highest uptake efficiency by transformants. The other negatively charged minerals, mica, and kaolinite as well as the overall positively charged mineral calcite showed almost the same transformation frequencies. The positively charged hematite displayed the next highest frequency and the positively charged goethite displayed the lowest transformation frequency. When we applied sub-lethal doses of antibiotics (Fig. 1, purple boxes) the relative trends persisted, and the transformation frequencies increased (Fig. 1). Plotting the transformation frequencies as a function of mineral surface the transformation frequencies for mica, kaolinite, calcite, and hematite appear to be positively correlated by the surface area. However, goethite and quartz, which have a similar surface area as mica, are outliers of the trend, which highlights that other parameters, additional to surface area, impact uptake frequency. In the following, we explore the relationship between both DNA-mineral binding affinity and bacteria-mineral interactions as determinants of the marked differences in transformation efficiency between the minerals.

**Fig 1.**
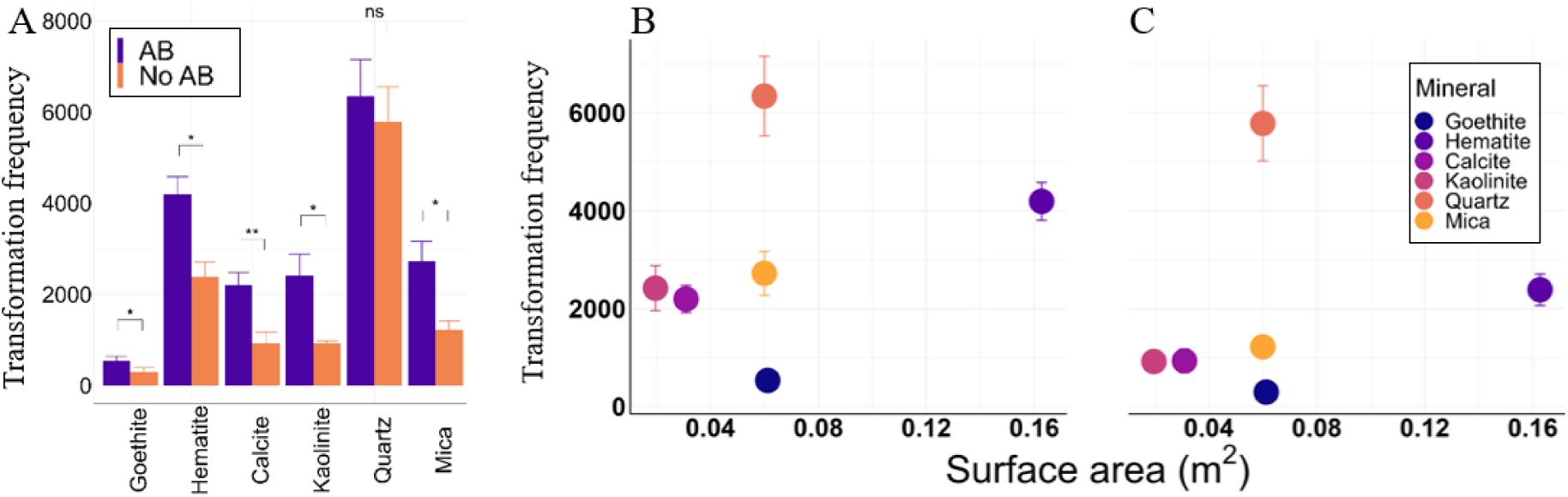
HGT frequency from mineral surfaces measured for *B. subtilis* with and without antibiotics. Data consist of three biological independent replicates. (A) Data are plotted in bar plots, where whiskers represent the standard deviation. For statistical analysis, we used Student t-test for independent samples to compare the two conditions (with and without antibiotics) for all minerals. * p<0.05., **p<0.01. Mineral concentrations used are listed in Methods. (B-C) Transformation frequency as a function of mineral surface area corresponding to the surface area used in the transformation experiments. A sublethal dose of antibiotics (AB) were added in B and not in C). Data is plotted in scatter plots, where the whiskers represent the standard deviation of the transformation frequency.

### Perspectives of DNA-mineral interactions for ARG transformation

As DNA itself is negatively charged, it is expected to be loosely adsorbed to the negatively charged surfaces, and thereby more available for bacterial uptake. Bacteria have the fastest transformation rate when encountering quartz. While quartz only has negative sites kaolinite, mica, and calcite have both negative and positive charged sites and goethite and hematite are dominated by positively charged sites. We know from AFM experiments that DNA preferably adsorbs to positively charged surfaces and charge dense sites and the adsorption strength of the DNA plays a role for the immobilization of an adsorbed molecules. For the mineral applied in this study we do observe distinct differences in how mobile the adsorbed DNA is. It is problematic to obtain AFM images of DNA mobility on kaolinite, hematite, and goethite. These minerals are nano to micro level particles and have topography corresponding to the topography of the DNA molecule itself and hence the DNA molecule is hard to resolve. In contrast mica, corundum and chlorite are close to being atomically flat and they both structurally and chemically resemble the silicates applied in this study; The mica serves as a model for the tetrahedral layer in kaolinite, the positively charged corundum is a model for octahedral clay layers as well as the octahedral clay edge sites.

The corundum surface retained the plasmid DNA while imaging in a tris buffer and in 10 mM MgCl2 solution. At the positively charged corundum surface, the DNA is adsorbed in a super coiled conformation highlighting a high amount of charge dense active adsorption sites [18]. Overlays of 3-7 frames in Figure 2. show in both cases that the DNA is adsorbed strong enough to be imaged, but the molecule is also wiggling as evident by the shifting red lines in the overlay images. The adsorption of plasmid DNA on chlorite shows the importance of charge dense adhesion sites for DNA mobility. Chlorite is a double layered clay mineral containing a brucite like layer (Mg(OH)_2_) and a mica like layer. The brucite like layer contrasts with mica positively charged. Overlay images of adsorbing plasmid DNA to chlorite in artificial seawater solution show that the molecule is immobile at the brucite edges and highly mobile over the areas of the mica-like surface. Chlorite data is from Sand et al [62]. Essentially the liquid cell AFM images (Figure-AFM1) show how local active site densities and binding strength influence the mobility of adsorbed DNA.

**Fig 2.**
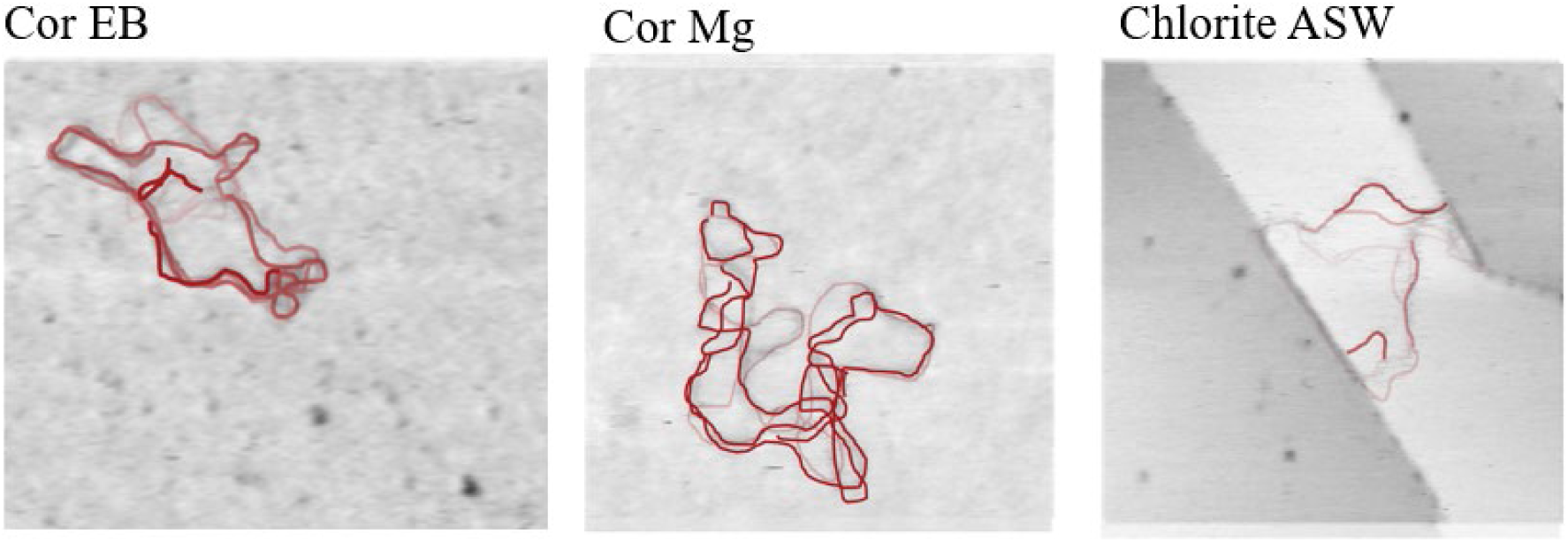
Maps of the mobile adsorption behavior of DNA molecules on different mineral surfaces. AFM images processed using MATLAB and overlayed to capture DNA movement through time. Corundum (cor) in EB-buffer (EB) (6 frames in 1;30 minutes), Cor in 10 mM MgCl_2_ (3 frames in 0:50 minutes) and Chlorite in artificial seawater (ASW) (7 frames in 1:42).

The active site density is a parameter that can influence how well a DNA molecule is adsorbed to a mineral surface. The reported values for transformation efficiencies are normalized to mineral surface area and a similar amount of DNA is added to each experiment. However, the active site density of each mineral varies independently of surface area. Usually, the active site density is measured via proton adsorption. DNA is a large molecule and sites available for protons and considering local changes in DNA adsorption visualized by AFM proton active sites is not a suitable indication of active sites for DNA adsorption. Instead, we used DNA adsorption data normalized to proton active site densities and surface area as a relative measure for active sites for DNA adsorption. Adsorption isotherms measured as adsorbed DNA concentration in ug/site density by Verma et al [3], show that goethite can adsorb approx. 3.5 times as much DNA pr surface area than hematite, and has a higher density of DNA-active-sites than any of the other minerals. In the transformation experiment 1 µg/ml DNA was adsorbed, leaving the active sites far from saturated. If we assume that a DNA-active-site is also an active site for interacting with bacteria there will be many sites available for bacterial interaction compared to the other mineral surfaces and hence a **lower probability for a fast encounter with a DNA molecule.**

### Perspectives of bacteria-mineral interactions

Another explanation for the variation in transformation efficiency could be caused bacteria-mineral interactions. We measured cell viability and cell integrity in suspension to understand if some of the minerals could cause cell death and hence influence the transfer frequencies observed in the HGT experiments above. Traditionally, cell viability in the presence of mineral is measured by adding the same amount of mineral mass to all experiments (Fig 3). Both plating and live/dead staining experiments from this setup show that viability is affected differently by the minerals added (Fig. 4 and Fig. 5). Following a 3 hr incubation of the bacteria with the respective minerals, we observed that a higher mineral concentration (1 mg/mL and 2 mg/mL) caused increased cell death (Fig. 3A and 3B)). Overall, the changes in viability were more severe for *B. subtilis* than for *A. baylyi*. The viability of *B. subtilis* was compromised by the minerals displaying positively charged surfaces: goethite, hematite, kaolinite, mica, and calcite. The viability of *A. baylyi* were only compromised in the presence of the iron oxides (hematite and goethite), although the live/dead staining did not show a significant number of dead cells for hematite (Fig 4. and Fig S3). Overall, the cells membrane integrity after exposure to all minerals for 3 hours using live/dead staining did confirm the trends observed in the viability experiments. The dead cell counts in the membrane integrity measurements are considered as minimum estimates of the sensitivity/specificity of propidium iodide in aggregates are quite low [63] and that dead cells that lyse after being dead is not visible by propidium iodide (PI) staining.

**Fig 3.**
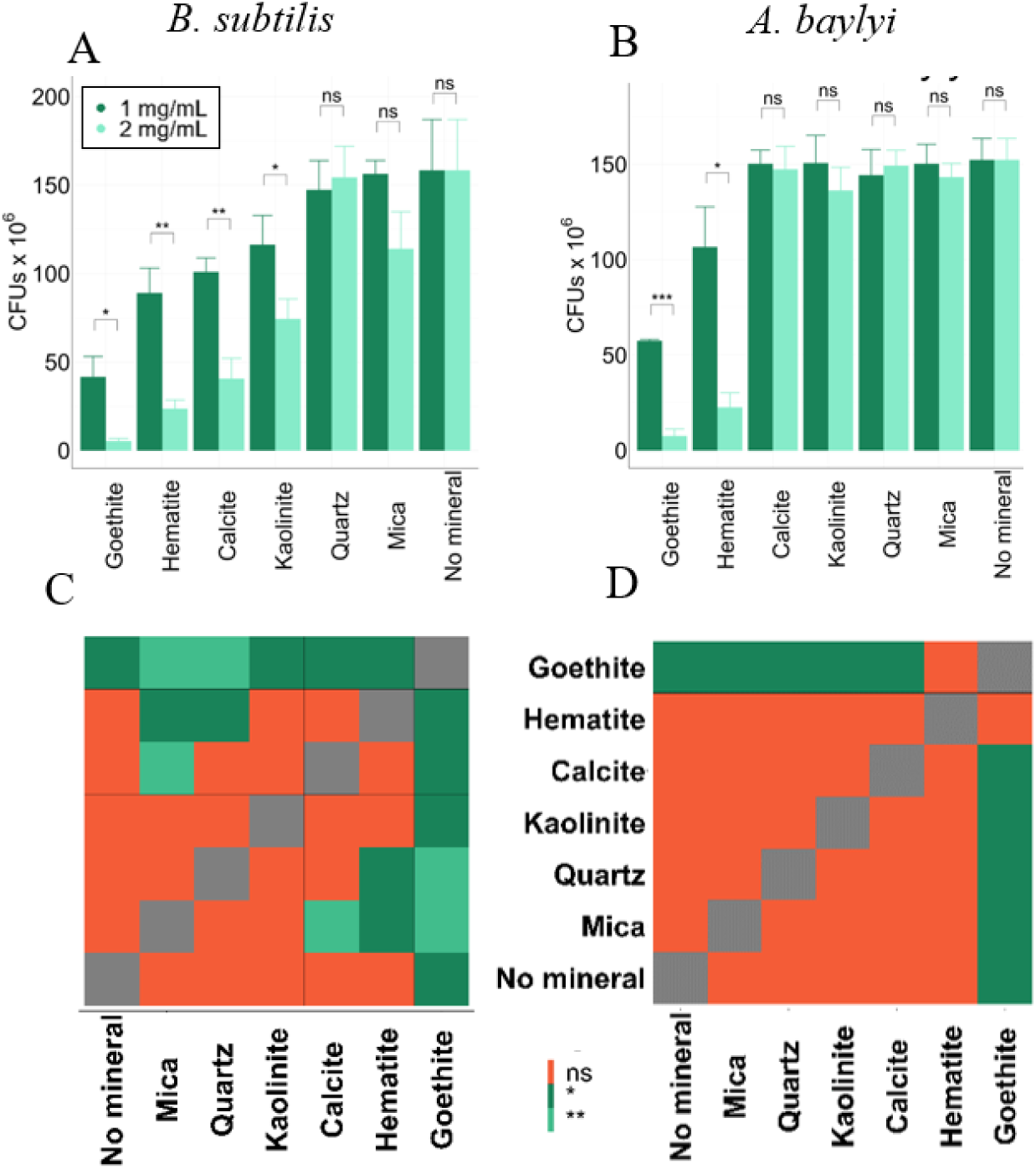
Effect of mineral concentration on cell viability of *B. subtilis* and *A. baylyi*. (A) Cell viability results for *B. subtilis* using two mineral concentrations (1 mg/mL in dark green and 2 mg/mL in bright green). (B) Cell viability results for *A. baylyi* using two mineral concentrations (1 mg/ml in dark green and 2 mg/mL in bright green). (A-B) *B. subtilis* and *A. baylyi* cells were exposed to the various minerals as indicated for a period of 3 h in TSB or LB medium, respectively. Post exposure, the cells were plated onto LB agar and incubated for 48 h. Colonies were counted and reported as CFU/mL. Mineral concentrations are the same used for the transformation experiments. Data consist of three biological independent replicates and are plotted in A-B using bar plots, where the whiskers represent the standard deviation of the CFUs. For statistical analysis, we used Student t-test for independent samples to compare the two conditions for all minerals. (C, D) Heatmap of statistical differences in cell viability among all minerals for *B. subtilis* and *A. baylyi*. Using the data from the cell viability experiments (Fig 3C) we performed comparisons among all minerals for the two bacterial species. We used Student t-test for independent samples to compare the different minerals and the FDR (False Discovery Rate) method for the correction of the p-values as we are performing multiple comparisons. Orange indicates no significant differences for the two minerals compared and the green tones indicate different significance levels. * p<0.05, **p<0.01. Exact numerical values for the P-values are listed in Supplementary.

**Fig. 4.**
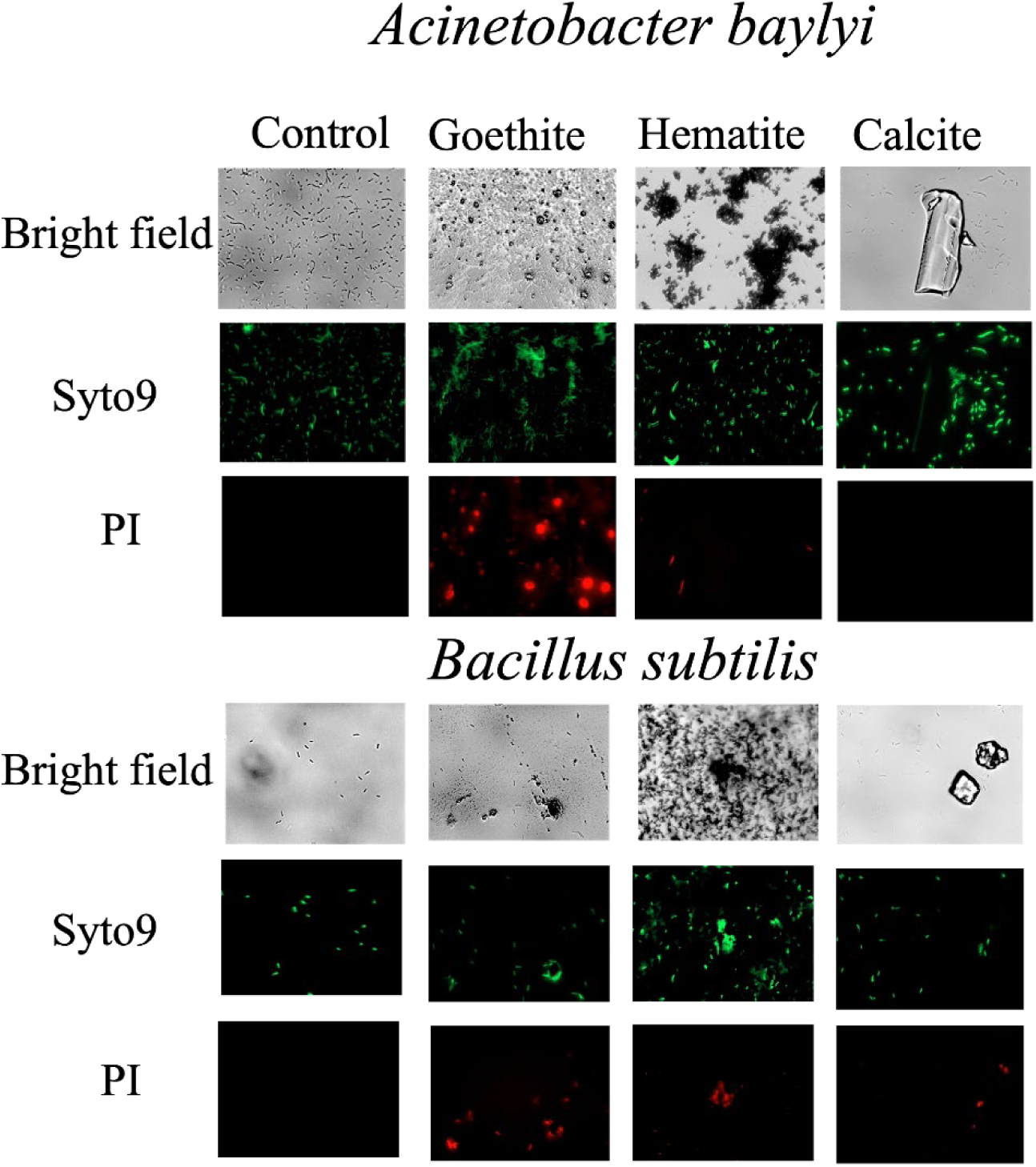
Cells membrane integrity. measured using fluorescent microscopy images of aliquots from the shaking culture sampled after 3 hr. (1 mg mineral). The cell permeable SYTO9 dye stains all DNA molecules with a green color including that in living cells. The cell impermeable propidium iodide (PI) stains both compromised cells and free DNA in a red color. Green free-swimming cells are observed in all images. The images shown are representative and all images in Fig S3.

**Fig 5.**
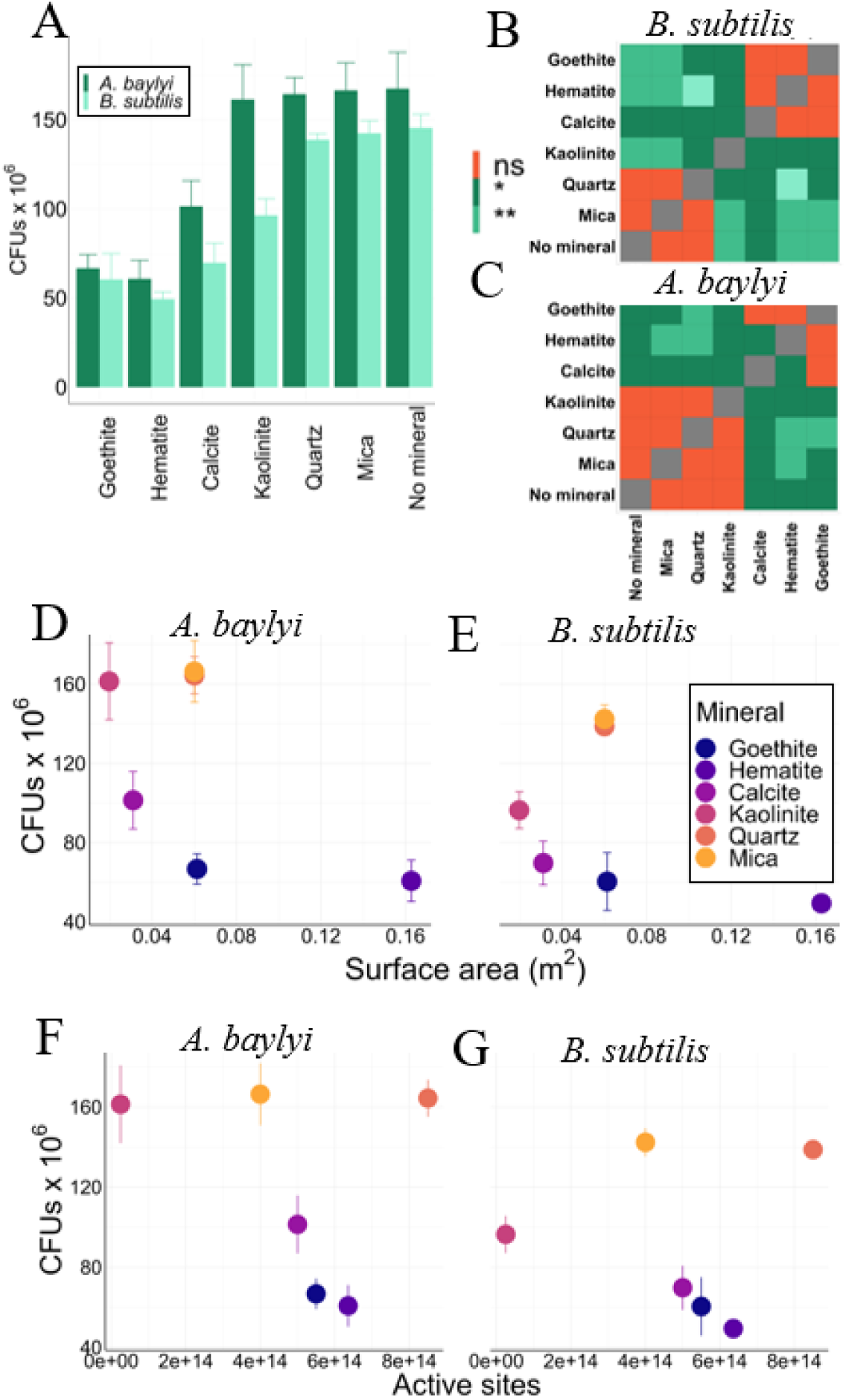
CFU as a function of. A) the mass of minerals as applied in the HGT experiments. The significance plots show the distinct groupings of the effect from the minerals. In general hematite and goethite (and calcite) have a similar influence and mica, quartz (and kaolinite) has a similar effect on viability. B-C) surface area (as applied in the HGT experiments) for *B. subtilis* and *A. baylyi*. Here Mica and quarts stand out with high cell counts compared to positively charged minerals with a similar surface area. Plotting against surface area D, E), there is a clear decrease in CFU with high surface area for the positively charged minerals. F-G) CFU as a function of available surface sites where for similar active site densities between the positive and negative charged minerals, the former show decreased CFUs and there is a marked inverse relationship site density and viability for the minerals with positively charged sites. Data consist of three biological replicates per experiment and are plotted in using bar plots (A) and scatter plots (D-G), where the whiskers represent the standard deviation of the CFUs. (C, D) Heatmap of statistical differences in cell viability among all minerals for *B. subtilis* and *A. baylyi*. Using the data from the cell viability experiments (A) we performed comparisons among all minerals for the two bacterial species. We used Student t-test for independent samples to compare the different minerals and the FDR (False Discovery Rate) method for the correction of the p-values as we are performing multiple comparisons. Orange indicates no significant differences for the two minerals compared and the green tones indicate different significance levels. * p<0.05, **p<0.01, ***p<0.001.

Measuring viability between the different minerals using the same mineral concentration does not account for the variation in mineral surface properties between the minerals. Reactivity of mineral surfaces are related to their surface charge, active site density and surface area which are all unrelated to mineral mass. In the HGT experiments we used a similar mineral surface area in all the experiments and hence different mineral masses. **When viability is measured using the same mineral mass as in the HGT** experiments, goethite, hematite, calcite and kaolinite influence viability (Fig. 5-A-C) (kaolinite does not appear to have an influence on viability for *A. baylyi*.) Significance heatmaps highlight the different grouping of viability between the minerals where for *B. subtilis* calcite, hematite and goethite have a similar effect on viability and for *A. baylyi* goethite and hematite have a similar effect (more in supporting info). If we plot the CFU counts (counted using the same mineral mass as added to the HGT experiments) as a function of the surface area of each of the minerals as present in the HGT experiments, there is a marked difference between the positively charged minerals and the negatively charged minerals. For a similar surface area, the minerals dominated by negative charges have lower viability (i.e. comparing quartz or mica with goethite). A similar effect is seen between kaolinite and calcite but more so with *A. baylyi*.

Low transformation rate of DNA from goethite surfaces has been argued to be caused by a goethite-killing effect where a strong association with the goethite surface punctures the bacterial cells. When we plot CFU as a function of mineral surface area (as applied in the HGT experiments) hematite shows the highest killing effect for both bacteria and for *B. subtilis* calcite has a similar effect. Additionally, viability for *both bacteria* is in general decreased when a positive surface is present (Fig. 5 D-G). With our approach of considering surface area and active site density it become apparent that viability decreases on surface when there is a strong bacteria-mineral attraction and even more so when there is a high number of active sites. At these minerals the bacteria is effectively adsorbed via numerous binding sites which potentially lead to cell lysis and/or increased stress response.

If we use DNA adsorption isotherms as a proxy for active site densities, bacteria on goethite will settle fast and the DNA will be many active sites apart compared to quartz and will both slow down bacterial mobility and decrease the transformation frequency. In contrast, quartz has a high number of active sites, but the interaction between bacteria and DNA is much weaker allowing a higher mobility of the bacteria and weaker adsorbed DNA. The optical images for quartz, calcite, mica, and kaolinite all showed a fair number of bacteria in the suspension and hence indicate some degree of freedom for the bacteria to mobilize (Fig S3).

### Perspectives of microbe-mineral interactions for ARG propagation

#### Biofilm formation

To test if the mineralogy plays a role for Arg dissemination we measured the rate of biofilm formation on pucks with deposited minerals, quantified protein content and protein structures using vibrational spectroscopy and subsequently tested the propagation of adsorbed DNA throughout the colonies during biofilm formation.

We used metabolic activity measurements in combination with SEM images to establish the extent of biofilm formation after 24 h incubations (Fig. 6). In general, metabolic activity and SEM images show an increased extent of biofilm activity when positively charged surfaces are present where Hematite shows the greatest extent followed by goethite, calcite, and finally mica, kaolinite and quartz showed the smallest extent of biofilm formation. In terms of metabolic activity, *B. subtilis* shows a larger variation in both metabolic activity and visual inspection of the extent of the biofilm formation than *A. baylyi*. Specifically, the OD405 values representing the amount of biofilm formation in control, goethite, hematite, calcite, kaolinite, quartz, and mica for *B. subtilis* were 0.88±0.49, 1.84±0.96, 2.45±0.4, 1.18±0.49, 0.64±0.19, 0.57±0.28 and 0.71±0.15 at 24h, respectively. The values for *A. baylyi* were 0.76±0.18, 1.45±0.23, 1.7±0.24, 1.1±0.22, 0.74±0.27, 0.69±0.26 and 0.7±0.22, respectively. The heatmaps in general show that goethite and hematite show distinct variation relative to the other minerals such as mica, quartz, kaolinite, and calcite (for *B. subtilis* calcite is distinct from quartz). Due to ingrained heterogeneity of biofilms, parts of the community can be in a low-metabolic or dormant state, and hence TTC measurements do not necessarily reflect the biovolume of the community but propose a minimum estimate of biofilm formation. A more nuanced view of biofilm extension is provided by the SEM images (Fig 6.), where it is evident that *A. baylyi* generally formed more patchy and thicker biofilm than *B. subtilis,* which instead formed more uniform and tangled thread-like biofilms that covered more surface area (explore relation to FTIR data).

**Fig 6.**
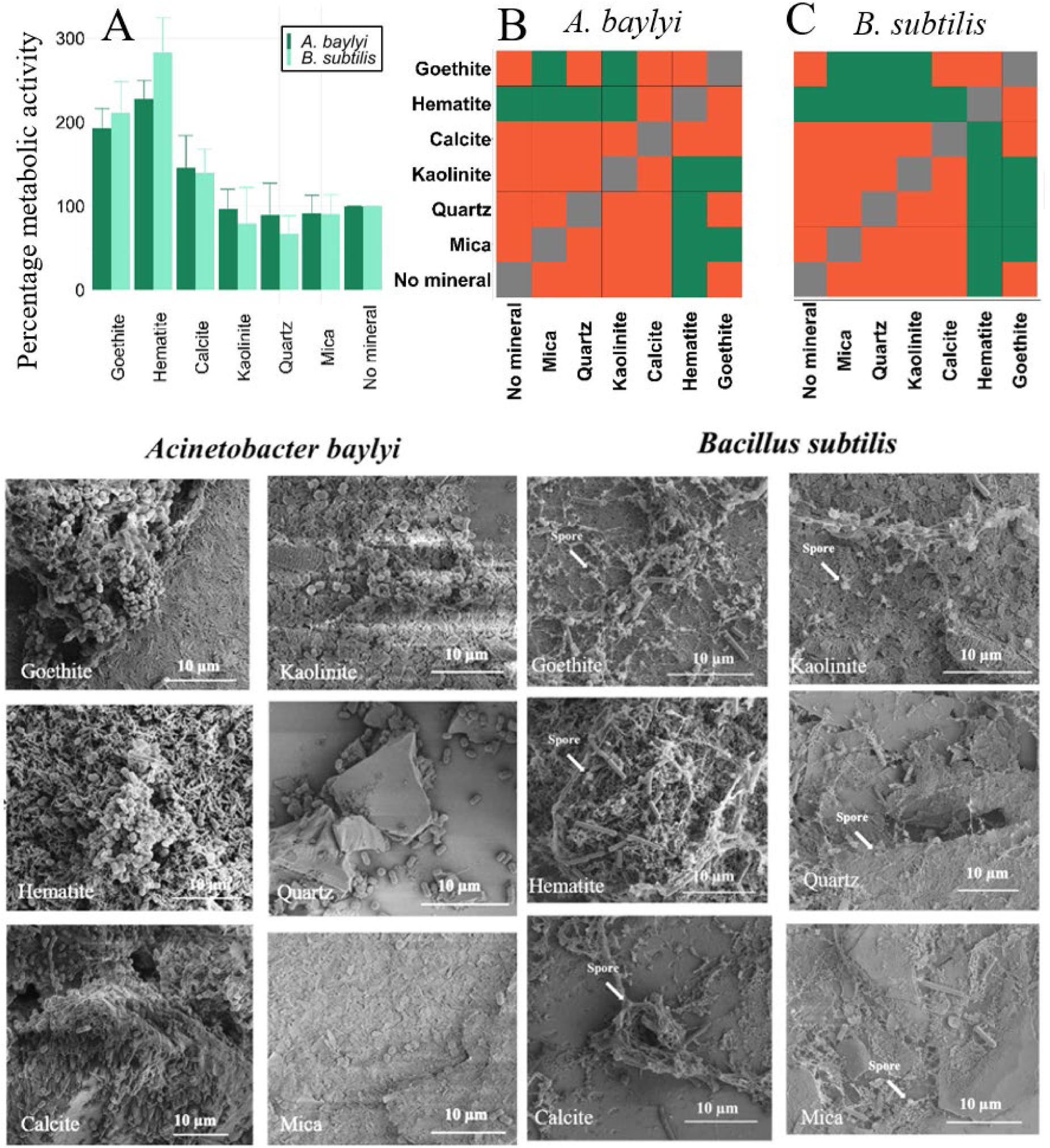
Bacterial metabolism is mineral dependent. A) The percentage biofilm formation based on TTC conversion was measured for *A. baylyi* and *B. subtilis* biofilms grown on glass coverslips coated with hematite, goethite, calcite, kaolinite, quartz, and mica, respectively, after 24 hours. TTC-converted formazan derivatives were measured at low frequency wavelength (405nm) and indicate metabolic activity. Controls for each bacterium were glass coverslips with no mineral coat and the OD405 value considered as 100% and all the OD405 values for biofilms on minerals were normalized based on that. Data consist of three biological independent replicates and are plotted in using a bar plot, where the whiskers represent the standard deviation of the metabolic activity. (B) Heatmap of statistical differences in bacterial metabolism among all minerals for *B. subtilis* and *A. baylyi*. We used Student t-test for independent samples to compare the different minerals and the FDR (False Discovery Rate) method for the correction of the p-values as we are performing multiple comparisons. Orange indicates no significant differences in the mean of the two minerals compared and green indicates * p<0.05. Exact numerical values for the P-values are listed in the Supplementary information. **D). Biomass and matrix production is mineral dependent.** With the application of scanning electron microscopy, the biofilm size and structure were evaluated after 24 h of incubation on various minerals for *A. baylyi* and *B. subtilis*. On kaolinite, quartz, and mica only a few bacteria were present, and no matrix could be found. In contrast, goethite, hematite, and calcite enabled multiple cells to attach and form a biofilm matrix. In addition, goethite and hematite also enabled the formation of spores integrated within the *B. subtilis* biofilm. Images are representative of multiple images acquired and solid lines indicate 10 µm.

### Effect of minerals on biofilm formation, structure and composition

The mineral surface characteristics of each mineral lead to quite distinct surface properties on the mineral pucks. We used AFM to measure the surface area of the mineral pucks. It was not possible to scan the entire surface of the puck so the surface area for the pucks is reported pr um^2^. The order of highest to lowest surface area are hematite, kaolinite, goethite, quartz, mica, and calcite (Figure 7A-B, Table S1). Using the active site density, we estimated the number of available sites pr unit area (Fig. 7C-D). From the surface area it is evident that calcite, goethite, and hematite show an increase in metabolic activity with surface area increase. For kaolinite, quartz and mica surface area do not appear to influence the metabolic activity. However, considering the active site densities we do see an overall correlation between metabolic activity. Quartz appears to have allowed metabolic activity considering its active site density which reflect that is less favorable to form biofilm on.

**Figure 7.**
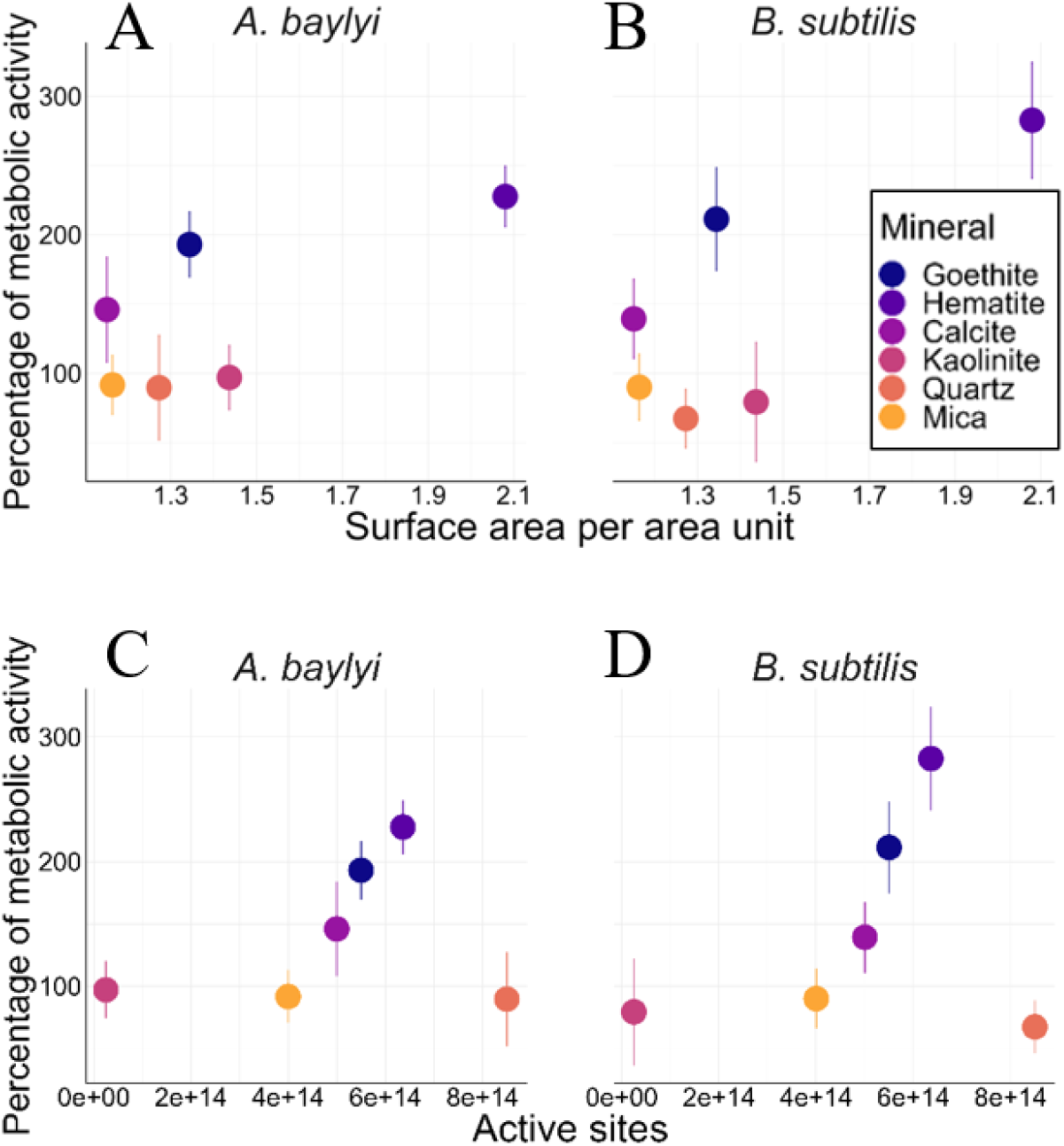
Percentage of metabolic activity plotted as a function of. A) Surface area and B) active site density for *A. baylyi* and *B. subtilis*. Data consist of three biological independent replicates and are plotted in using a scatter plots, where the whiskers represent the standard deviation of the percentage of metabolic activity.

Biofilm formation and development, from attachment to dispersal, is affected by various physiochemical conditions. Substrate topography is a key determinant of biofilm formation [60] [64-66] that with reason can be expected to vary among different types of minerals. With the use of gold substrate terminated with self-assembled monolayers of various chemicals, we also previously found that positively charged and hydrophobic particles support biofilm formation, while negative surface charge and hydrophilic particles resist bacterial communities to congregate [67]. The strong attraction to positive charged surfaces and vice versa repulsion to negative charged particles, can potentially be explained by the general net negative charge of bacterial cells [68, 69]. This trend is also reflected in the particles tested here, where the positive hematite and goethite are the substrates with highest metabolic activity and with evident aggregation (Fig. 6). In suspension, calcite, hematite, and goethite were also associated with a relatively large number of death cells (Fig.4 and Fig 5)). In principle, dead cells can be perceived as biofilm matrix, and cell death as part of a coordinated biofilm development [70]. Perhaps this behavior is induced by positively charged minerals.

Although cell survival assays demonstrated a killing effect of calcite, goethite and hematite, still high amount of metabolic activity is recorded and biofilm buildup is confirmed with SEM. Overall, the bacteria show high willingness towards adsorption to these minerals. In suspension, *A. baylyi* did show less cell death compared to b. subtilis (as a function of surface area Fig. 5) which could be due to the different biofilm architecture. For *A. baylyi* the biofilm developed in patches whereas the b. subtilis showed more extended coverage. The buildup seen for biofilms for *A. baylyi* would decrease the contact with the mineral surface pointing toward the surface structure or bacteria-mineral interaction for playing a role for cell death. Having a surface that compromise cell integrity would induce biofilm formation as a protective stress response, as also observed for other antimicrobials [71]. In this case, the biofilm matrix could be supported by the positively charged surface of the mineral due to attraction of lysate, which mainly is constituted by negatively charged cell wall material, DNA and metabolites [72]. This attraction of lysate could then function as matrix but also as a nutrient boost for surviving cells and escalate metabolic activity [73, 74].

## DNA-BACTERIA-MINERAL

Propagation of ARG in the environment is dependent on how well a transfer event can propagate throughout the colony. Here both transformation, conjugation and cell division add to the propagation. We absorbed a similar concentration of DNA to mineral pucks and allowed biofilms to form. Transfer efficiencies were tracked after 3 hours and 24 hours with and without sublethal levels of antibiotics. In general, the transformation efficiencies seen after 3 hours (Fig. 8A) follow the trend as seen for uptake frequencies from the suspended minerals and an amplification with added antibiotics. Significance heatmaps (Fig 8B) show that, at sublethal antibiotics) quartz that the highest frequency followed by hematite, then kaolinite, mica and calcite impacting efficiency equally, and finally goethite again displaying the lowest frequency. The surface area of the pucks may have slight positive correlation with transformation efficiency excluding quartz and goethite and likewise considering active sites pr surface area (Fig 8A-D).

**Fig 8.**
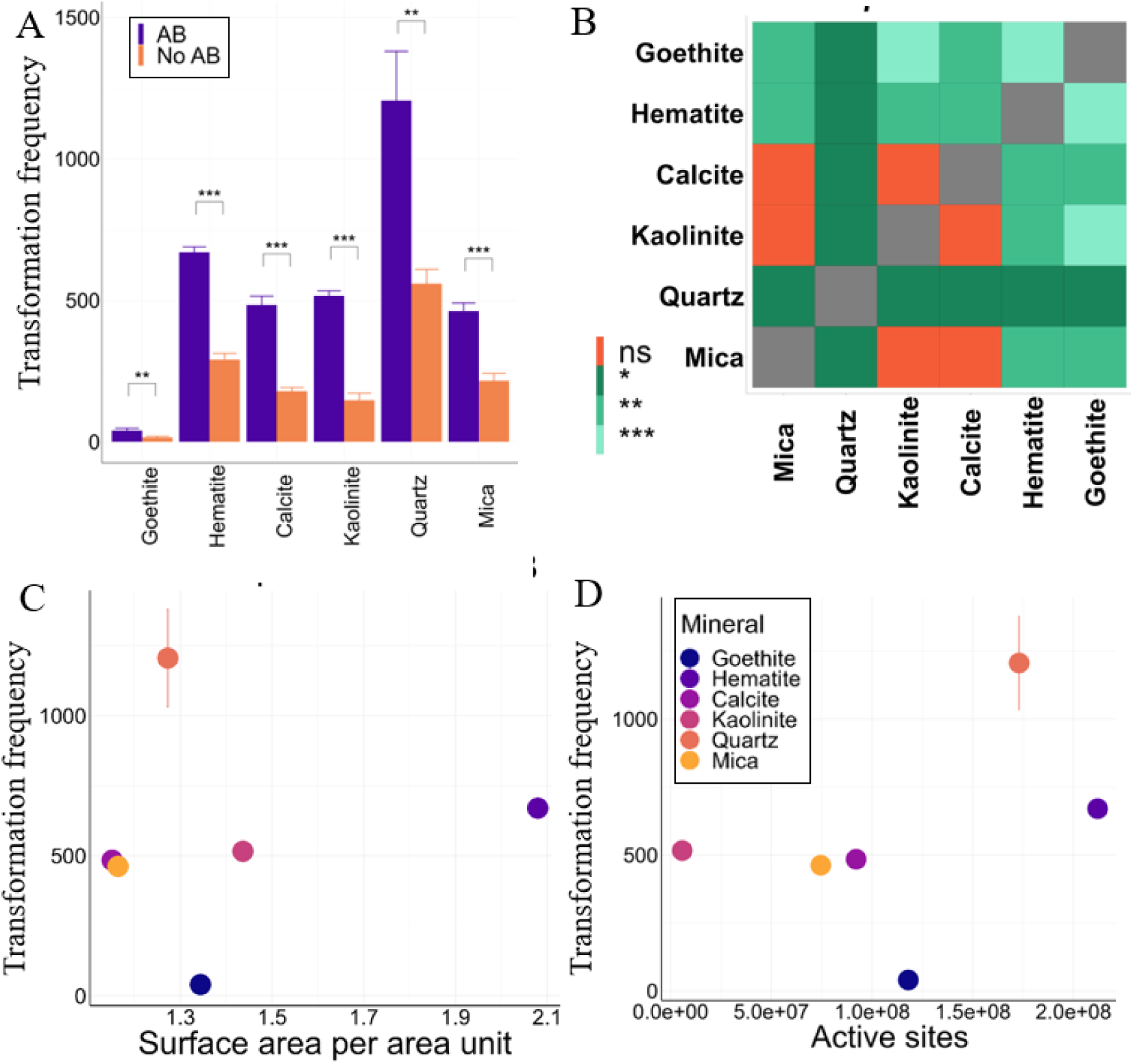
A) **HGT from mineral pucks after 3 hr.** with and without AB. B-D with sublethal doses of AB. B) significance heatmaps, C) transformation frequencies vs surface area, D) transformation frequencies vs active sites pr area. D and E are only shown for +AB (-AB is in the supporting Fig S6).

After 24 hours of biofilm formation the transformation frequencies are high for the minerals that also enhanced biofilm formation which appear inverse from the observations seen in the transformation experiments (Fig 9.). The transformation frequencies are counted as both events from cell division and transformation. As seen for the biofilm formation experiments, the higher the surface area and the active site density the faster propagation of the trait. According to the heatmap, there are only significant differences when comparing Quartz with Kaolinite and when compared with Calcite and the number of transformation frequencies following propagation of the DNA transferred from the minerals tend to follow a similar trend for all the minerals, maybe because of the biofilm formation.

**Figure 9.**
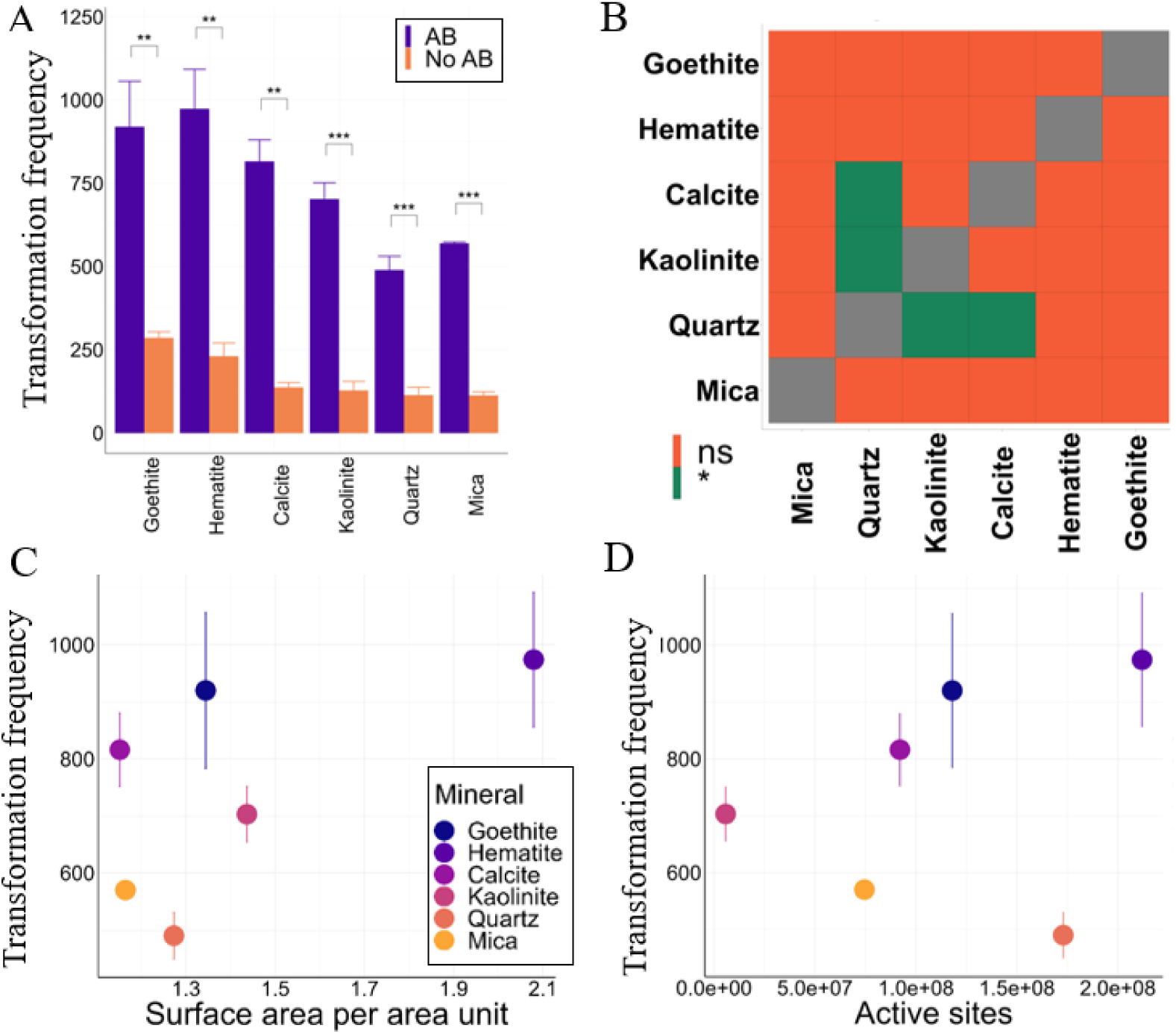
A) **HGT from mineral pucks after 24 hr**. with and without AB. B-D with AB. B) significance heatmaps, C) transformation frequencies vs surface area, D) transformation frequencies vs active sites pr area.

## DISCUSSION AND IMPLICATION

Transformation of Mineral adsorbed DNA to the competent bacteria B. subtilis do, as reported by other studies, show variation in transformation frequency with mineral surface. The transformation in our case was enhanced when we induced a selective pressure by adding sub-lethal antibiotic doses. Adding antibiotics can enhance the growth rate of bacteria harboring resistant plasmids, leading to a competitive advantage over non-resistant counterparts within mixed bacterial populations [75, 76]. Moreover, sub-lethal antibiotic concentrations foster heightened rates of HGT, further facilitating the dissemination of resistance genes [77]. Regardless the mechanism, the relative frequencies of uptake for the different minerals as seen without selective pressure persists.

From our data, we find the main driver for the control minerals exerts on both, transfer efficiency, bacteria viability, biofilm formation or ARg propagation in a colony to be the surface properties: surface charge, surface area, active site densities and likely the strength of interaction between the DNA and the mineral surfaces. The mechanism driving the influence from mineral surface properties could both be driven by biologic processes as well as related to interfacial geochemical reactions. Regardless, our data show that mineral surface properties influence a wide range of bacterial processes and could be considered as a tool to guide mitigation strategies for preventing further ARg propagation in our environments.

In the transformation experiments an increased surface area appear to play a role for enhancing the transformation frequency except for quartz and hematite. <u>Considering the interfacial geochemical</u> <u>interactions</u>, quartz have a high transformation frequency and is also the mineral with only negative surface charge. The negative surface charge can weaken the DNA-mineral bond, potentially making uptake easier for the bacteria. The negative surface can also cause a weak interaction with the bacteria and enhancing bacterial mobility in the aqueous phase relative to a highly positively charged mineral. In the case of a positively charge surface as seen for calcite and hematite, the stronger electrostatic interaction with the bacteria can enhance the driving force for interacting with the mineral surface and hence enhance the probability of encountering the adsorbed DNA. Calcite has different types of binding sites and a lower site density than hematite which would make the hematite surface more attractive toward bacterial interaction.

Considering the viability experiments, when applied in the same concentrations as in the HGT experiments (same surface area for all minerals) both calcite, hematite and goethite decrease bacterial viability to a similar extent (*for B. subtilis*). The reason for low uptake frequency for goethite can be a question of faster killing kinetics or because of a strong interaction between goethite and the bacteria preventing mobility and hence a strong decrease in probability of encountering the adsorbed and strongly bound DNA. Accounting for active surface area and available surface sites (Fig. 5) it is evident that there is a distinction between the positively and negatively charged minerals where a high density of active sites has a negative influence on viability if the mineral is dominated by positive charges. However, there is no correlation between transformation efficiency and viability for the minerals with a high density of positive sites as goethite show a low transformation frequency and hematite a high frequency. The variation can be explained by adsorption capacity. When measured against active site density goethite has a high adsorption capacity for DNA and likely also for signaling molecules Hematite has a lower adsorption capacity and displays a higher transfer efficiency.

During biofilm formation the metabolic activity and the surface buildup is again higher for hematite and goethite. Here, immobilized on a mineral puck the calcite effects the biofilm in a similar manner as the quartz, mica, and kaolinite. The metabolic activity appears to be correlated with available surface sites, again indicating that higher charge density on the mineral surfaces enhances biofilm formation as well as the metabolic activity.

Adsorbing DNA to mineral pucks and allowing the DNA to propagate over the course of 3 and 24 hr the minerals with positive surface sites has a low transfer efficiency after 3 hours (similar to the transformation experiments in solution): however, after 24 hours the trends are reversed and we observe a marked higher propagation efficiency for the positively charged minerals when the biofilms are allowed time to establish.

<u>From a biological point of view</u>, the mechanism behind the bacterial response causing the influence of the mineral surface can be multifold and not explored to a great extent here. We argue that it can be a multitude of mechanism acting in both concert and counter active. i.e.: Signaling molecules are adsorbed causing loss of competence, although our bacteria are competent when we start the experiments. Adsorption of competency genes are expected to follow the same general trends as DNA and again, there is a high number of available sites for adsorption of competency molecules on goethite relative to any of the other minerals. However, the bacteria applied here are competent from onset and we expect any effects of adsorption of competency molecules to play a minor role.

- Release of ROS from goethite, hematite, and calcite that affect bacterial survival in lethal doses. Once killed the bacteria can act as food and as anchor points causing a surge of cell growth and biofilm formation, also enhancing cell division and uptake. Induction of oxidative stress in sub-lethal doses inside the cells influences a cascade of serially different regulatory changes, by changing in the protein expressions patterns. For instance:

- inducing the upregulation of stress response proteins, such as chaperones and heat shock proteins, to counteract the adverse effects of mineral-induced stressors.
- By triggering the expression of membrane-bound transport proteins involved in the uptake of essential nutrients and ARG or the expulsion of toxic compounds.
- modulating the expression of virulence factors, such as adhesins and toxins, potentially influencing the pathogenicity or attachment of the bacterial cells.
- Overexpressing the quorum sensing and other regulatory gene expression can lead bacteria to form biofilms on mineral surfaces.

Once taken up by these naturally competent bacteria, plasmid DNA can be horizontally transferred to many other bacterial species (not only the naturally competent ones). We consider mineral facilitated gene transfer of extracellular DNA as the first step for further dissemination in bacterial populations in the environment. Regardless the mechanisms driving the reactions, several of the mechanisms can act in concert guided by the mineral surface properties. Consequently, the positively charged mineral can be considered as hot sports for both DNA preservation, biofilm formation and enhanced propagation of transformed extracellular DNA. The DNA-mineral bond strength may be determining for initial uptake frequency, but given enough time, the DNA are taken up. Mitigation wise it is hard to combat both DNA preservation (enhanced at positively charged surfaces) and biofilm formation with its subsequent opportune influences of trait propagation. Hence degradation may be the best approach for preventing propagation via mineral facilitated transfer of extracellular DNA.

## CONCLUSIONS

The transfer frequency of DNA adsorbed to mineral surfaces in to B. subtilis changes with mineral surface properties, and so did viability. Instead of measuring viability of *b. subtilis* and *A. baylyi* by comparing between mineral concentrations (mg/mL) we measured viability using a mineral concentration that provided similar surface area of all of the minerals. Here it become evident that cell viability in general is decreased at minerals displaying a high density of positively charged sites. However, the viability does not correlate with transformation efficiency and hence other parameters than viability play a role. We here argue that those could be related to the DNA-mineral bond and/or general adsorption capacity for signaling molecules. The rate of biofilm formation for both *b. subtilis* and *A. baylyi* are also affected by mineral surface properties in particular surface area and active site density. Here the surfaces with high viability show enhanced biofilm formation.

We show that mineral surface characteristics well can drive initial interactions during biofilm formation and transformation of mineral adsorbed DNA. When the biofilm develops, the control on trait propagation as exerted by the mineral surface diminishes and all the surfaces are able to considerably propagate ARg.

We argue that consideration of surface properties can be utilized in mitigation strategies for preventing the propagation of ARg, however, it is concerning, in the light of mitigation, that once the biofilm develops the traits will be propagated regardless of the surface properties.

## ACKNOWLEDGEMENTS

We wish to thank Milda Pucetaite and the Microscopy Facility at Department of Biology, Lund University for FTIR guidance and access to the Vibrational Spectroscopy lab. We also acknowledge Klaus Qvortrup and the Core Facility for Integrated Microscopy, Faculty of Health and Medical Sciences, University of Copenhagen for help with the SEM images. The work presented here is supported by the VILLUM FONDEN grant 00025352 (KKS, SH, TV), by the Carlsberg Foundation grant CF21-0311 (KKS and CM), and by the Danish National Research Foundation, grant DNRF174, Centre for Ancient Environmental Genomics (CAEG) (PA). OA is grateful to the European Union’s Horizon 2020 research and innovation programme under the Marie Sokolowski-Curie grant agreement No. 892889. We also acknowledge the Slovenian research and innovation agency which is financing the national research program P4-0116 (IM)

## AUTHOR CONTRIBUTIONS

KKS, SH, MB, IM, designed the study; SH performed bacterial experiments; MH and SH performed confocal microscopy, MH made statical analyses of the confocal microscopy data. TV helped with initial DNA adsorption experiments, CCM, PA, KKS performed and analyzed atomic force microscopy. KKS, SH and CCM interpreted the data, KKS, SH, CCM, OA, PA, TV, MH, IM and MB were involved in writing the manuscript.

## METHOD SECTION

### Material and methods

#### Bacterial strains and culture conditions

Soil-habitat Gram-positive *B. subtilis* 168 and Gram-negative *A. baylyi* BD413 were used in this work. The strains were first cultivated in separate pure cultures overnight on Tryptic soy agar (TSA; Merck, Germany) at 37°C. Single colonies of *B. subtilis* and *A. baylyi* were initially grown overnight in 25 ml of Tryptic soy broth (TSB; Merck, Germany) for *B. subtilis* and Loria berthany (LB; Merck, Germany) for *A. baylyi*. These initial cultures were incubated at 37°C with shaking at 180 rpm for 18 h corresponding to ∼10^8^ cells. The cells were centrifuged at 5000 rpm at 4°C for 10 mins and resuspended in prechilled LB containing 15% glycerol, aliquoted and stored at -80°C until further use.

To prepare competent *B. subtilis* 168 cells, SPIZIZEN (SP) media was used which is composed of 14 g/L K_2_HPO_4_, 6 g/L KH_2_PO_4_, 1 g/L Trisodium citrate, 40 ml Glucose 50% (w/v), 20 ml Casein hydrolysate 5% (w/v), 10 ml Tryptophan 5 mg/ml, 500 µl Ammonium ferric citrate 22 mg/ml, 5 ml Potassium glutamate 40% (w/v), 3 ml MGSO_4_ 1M and 1ml spectinomycin 100 mg/ml. Single colonies of *B. subtilis* 168 were grown overnight in 5 ml SP media at 37 °C and 180 rpm. The 5 ml overnight culture was then transferred to 250 ml of fresh SP and incubated at 37°C under shaking at 180 rpm (for ∼4.5 hr). Cultures were harvested at the late-exponential phase by centrifugation at 5000 rpm for 30 mins. The bacterial concentration was adjusted to an OD600=0.6, corresponding to approximately 10^8^ CFU/ml.

### Horizontal gene transfer experiments

The minerals were weighed according to their surface area as listed in Table X to have the final weight with same surface area. Each mineral was disaggregated using a sonicator for 30 mins and autoclaved at 121^0^C for 20 minutes to remove any microbial contamination and resuspended in 1 mL sterile 150 mM NaCl at pH 7. Minerals were then mixed with the appropriate concentration of pEM1069 plasmid (as derived from isotherms results in our previous study) in a way that the DNA adsorbed by the minerals was ∼ 1 ng/µL. The mineral-DNA suspensions were shaken overnight at room temperature at 70 rpm and centrifuged at 5000 rpm for 30 mins. The residues were washed with 1 mL sterile saline until no free DNA was detected in the supernatant (estimated by measuring absorbance at 260 nm). The mineral-DNA pellets were then used for transformation experiments.

For horizontal gene transfer experiments, the “Paris method” with SP minimal media was conducted (Harwood, C. R., and Cutting, S. M. (1990)). The competent cells were thawed on ice and washed once with sterile saline. The cells were then resuspended in SP media, added to the mineral-DNA pellets, and incubated for 2 hr at 37^0^C under shaking at 180 rpm. Post incubation, the cells were plated on LB agar containing 100 µg/ml spectinomycin and incubated at 37^0^C for 24 h to screen for transformants. Controls were:

- *B. subtilis* cells exposed to the minerals in the absence of DNA and plated both on LB with spectinomycin as well as without spectinomycin.

- *B. subtilis* cells with plasmid in the absence of minerals and plated on LB plus spectinomycin. The transformation efficiency per µg DNA was calculated as follow:

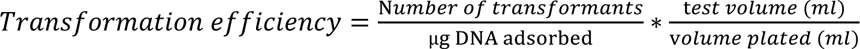

The transformation frequency is normalized by mineral surface area and a similar amount of DNA is adsorbed to all the mineral surfaces (method section).

### AFM image overlay

AFM images were exported from Igor Pro software and processed in MATLAB to obtain the negatives of the images. Then the images were overlayed using the features of the mineral surface as a guide and marking the DNA strands to track their movement throughout time.

### Bacteria survival

To test the bacterial survival after being in contact with mineral surfaces, 200 µl of bacterial suspension (OD600=0.1) was incubated in 100 µl of two different concentrations of minerals (final concentrations of 1 and 2 mg/ml) and 1 ml TSB or LB media in a 5 ml sterile centrifuge tube for *B. subtilis* and *A. baylyi*, respectively. After shaking at 180 rpm for 3h at 30 °C, bacteria were washed in saline and decimal serially dilutions were made. Dilutions were then immediately spread on LB agar plates and incubated at 30 °C for CFU enumeration.

### Cell membrane integrity

The membrane integrity of *A. baylyi* and *B. subtilis* after 3h exposure to minerals was assessed by the LIVE/DEAD Bac Light Viability Kit, L7012 (Fisher scientific) according to the manufacturer’s instruction. The kit includes two DNA-binding stains: the green fluorescent stain, SYTO9, and the red fluorescent stain, propidium iodide (PI). For fluorescent imaging, we used an Axio Inverted Observer Microscope Z1 (Carl Zeiss Inc.) equipped with an Axiocam 503 mono camera and a N-Achroplan 63x/0.85 objective. After 20 min of staining in 10uM Syto9 and 60uM Propidium iodide (Filmtracer LIVE/DEAD viability kit, Thermo Fisher), samples were placed between a #1.5 cover glass (Thorlabs, CG15KH1) and an agar pad to reduce desiccation. Stained cells were illuminated with reflected light from a HXP 120C light source and filter set 44 FITC (475/40, 530/50) and filter set 43 DsRed (545/25, 605/70), respectively. Bacteria with intact cell membranes were stained fluorescent green and alive, while those with red fluorescent are membrane damaged and dead.

### Bacterial metabolic activity

To test the bacterial metabolic activity on mineral surfaces, mineral-coated 13 mm round glass coverslips were covered by mineral suspensions. Briefly, 0.4 ml of the ultrasound mineral suspension (1 mg/ml) were pipetted onto the coverslips and the coverslips were then boiled at 120°C for 30 min. The coverslips were rinsed using deionized water and dried at 45°C following by sterilization by autoclaving for 20 min at 121°C. Sterile mineral coated coverslips were placed in sterile 48-well flat-bottom polystyrene plates (Costar, Corning Incorporated, Corning, NY) and supplemented with 1 ml of bacterial suspension. The plates were incubated statically at 28 °C for 24h. Finally, Coverslips were immersed in sterile deionized water twice to eliminate loosely adhered bacterial cells. The metabolic activity of attached cells to mineral-coated coverslips was determined using triphenyl tetrazolium chloride (TTC, sigma aldrich). 0.5% TTC solutions was prepared in distilled water and sterilized by 0.22 μm cellulose acetate filters. Coverslips were transferred to a new sterile 48-well plate. 200 μL of TTC solution and 800 μl of TSB or LB were added to all the wells for *B. subtilis* and *A. baylyi*, respectively. Plates were incubated in the dark for 24h at 37 °C. After the incubation period, 200 μl of the well contents were moved to a new 96-well flat-bottomed microplate (Costar, Corning Incorporated, Corning, NY) and the absorbance was measured at 405 nm.

### Scanning electron microscopy

Morphology of biofilms formed on mineral coated coverslips was visualized using scanning electron microscopy (SEM). 24h biofilms were washed twice in fresh NaCl solution (9 g/l in water), and SEM samples were prepared following the classical steps: samples were chemically fixed by immersing overnight in a 4% glutaraldehyde (w/v in 0.1 PBS) solution, dehydrated by immersing in several ethanol solutions (from 20% (v/v) in water to 100%). Samples were dried using a critical point dryer (Leica (Balzer) CPD030). Finally, coverslips got sputter-coated with gold (Leica Coater ACE 200) at room temperature.

Examination was done at several magnifications (from 5000× to 100,000×) using a versatile high-resolution dual beam SEM (FEI Quanta 3D FEG) with high vacuum mode, Everhart-Thornley Detector (ETD). For each sample, several places were focused, and a series of images was taken by systematically displacing the samples under the camera.

### Data analysis

Data were analyzed in R. The statistical analysis was performed using the rstatix package.

