## Supporting for "Reconciling the importance of minerals for propagation of antibiotic resistance genes in the environment"

**SUPPORTING INFORMATION**

**Table S1.** Surface area (m2/wt%), Active sites on the total surface area (per nm2) and active sites on the total surface area (sites/cm2) were calculated for all the minerals in suspensions. Active sites (1/cm2) and surface area (1/cm2) were calculated for each mineral on pucks. Overall surface charge is presented for each mineral.

|  | **Suspension** | | | **Pucks** | | |  |
| --- | --- | --- | --- | --- | --- | --- | --- |
|  | Surface area as used for HGT | Active sites on the total surface area used for HGT | Active sites on the total surface area used for HGT | Active site density | Puck surface area | Active sites available pr unit area | Overall surface charge |
|  | m2/wt% | pr nm2 | sites/cm2 | 1/cm2 | 1/cm2 |  |  |
| **Mica** | 9.01E-04 | 5.73E-03 | 5.73E+11 | 4.00E+14 | 1.86E-07 | 7.45E+07 | -/+ |
| **Quartz** | 9.01E-04 | 4.96E-03 | 4.96E+11 | 8.50E+14 | 2.04E-07 | 1.73E+08 | - |
| **Kaolinite** | 9.01E-04 | 2.25E-04 | 2.25E+10 | 2.50E+13 | 2.30E-07 | 5.75E+06 | + , - |
| **Calcite** | 9.01E-04 | 3.60E-03 | 3.6E+11 | 5.00E+14 | 1.84E-07 | 9.21E+07 | + |
| **Hematite** | 9.01E-04 | 4.51E-03 | 4.51E+11 | 6.36E+14 | 3.33E-07 | 2.12E+08 | + |
| **Goethite** | 9.01E-04 | 7.66E-03 | 7.66E+11 | 5.50E+14 | 2.15E-07 | 1.18E+08 | + |

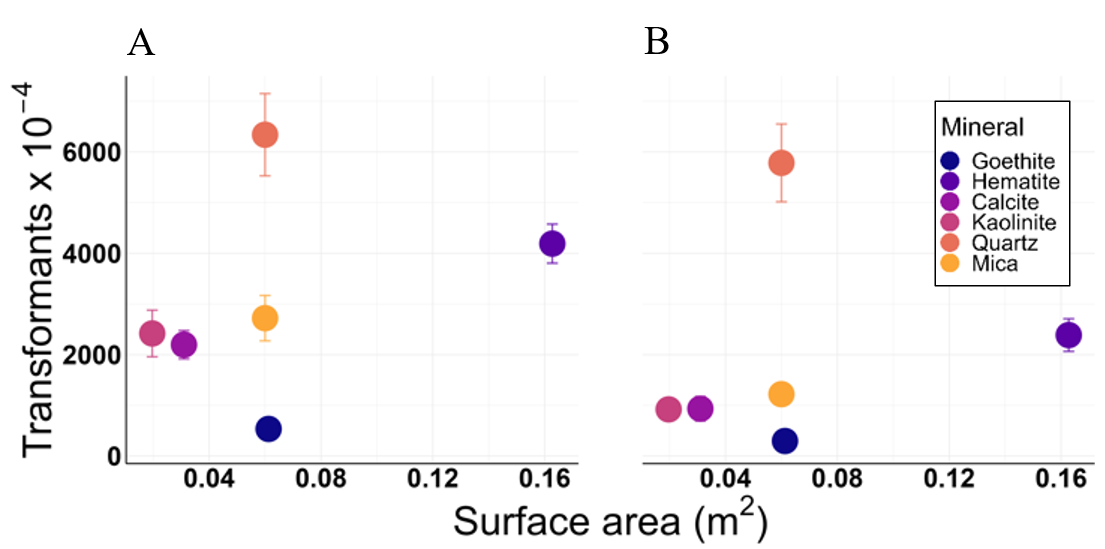

**Fig S1. Transformation efficiency vs. surface area** of the minerals the DNA is adsorbed to. No correlation between uptake efficiency and surface area.

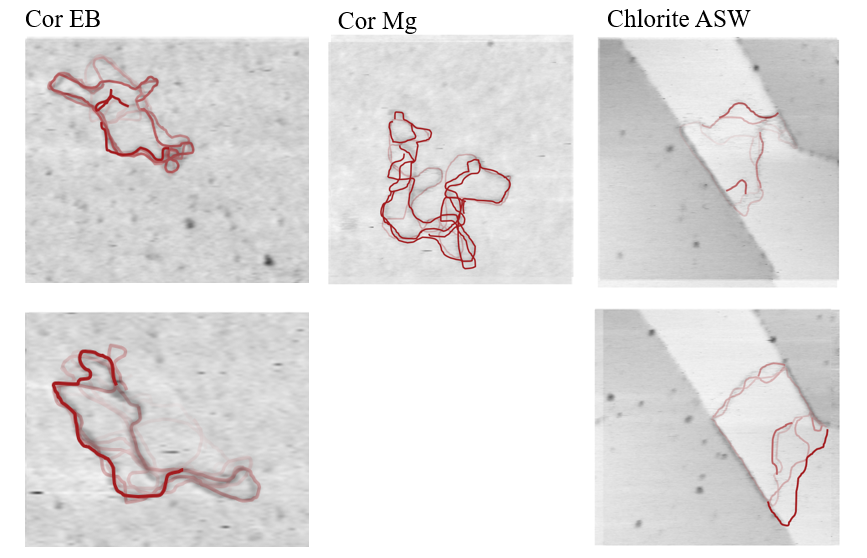

**Fig S2**. Fast scanning AFM imaging in liquid of plasmid DNA moving on the surfaces of A,B) Corundum in EB buffer, C) Corundum in 10 mM MgCl2 and in D,E) Chlorite in artificial seawater. The plasmids are about 200 nm in width.

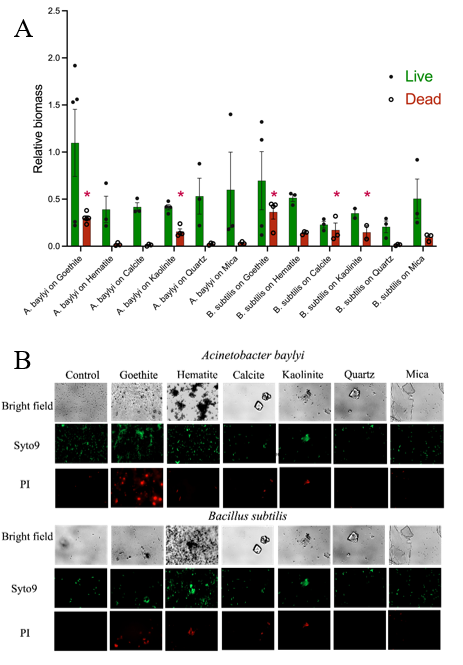

**Fig S3. Fluorescent microscopy images of aliquots from the shaking culture sampled after 3 hr**. (**1 mg mineral**). A) *B. subtilis* are more affected by *A. baylyi* and there are a significant number of dead biomasses for goethite, hematite, calcite, kaolinite and mica. For *A. baylyi*, goethite and kaolinite influence bacteria viability. Because of a variance, there is no significant difference between the biomass of living cells on any of the minerals, or between species (Brown-Forsythe and Welch ANOVA w. Dunnetts multiple comparison). B) The first row for each bacterial cells, shows Bright field images mineral and bacterial cells without fluorescent dye. The second row shows all alive cells that are stained by the cell permeable SYTO9 dye. The last row shows that the cell impermeable propidium iodide stains both compromised cells and free DNA in a red color. These images showed the killing effects of goethite for both bacterial cells. A slight influence of hematite for two strains and calcite, kaolinite, and mica for *B. subtills* was observed. While quartz was not able to affect cell membranes.

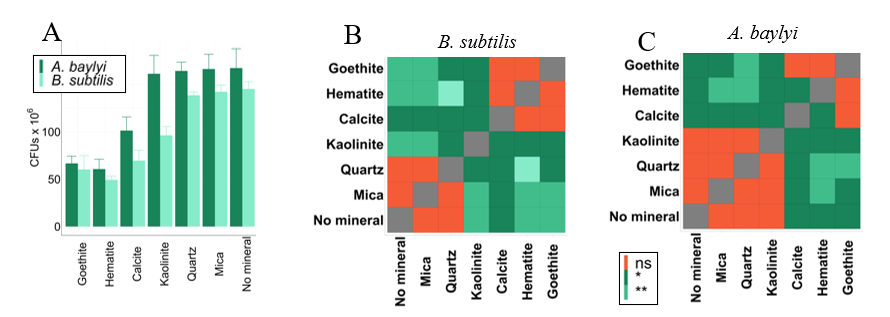

**Fig S4.** **Cell Viability**. Fig. 5 is restated here for clarity.

When we track cell viability using the same concentrations as for the HGT experiments in Fig S4. we see some minerals have a stronger effect on the survival of the bacterial cells, especially those with positive charges in their surface (Fig S4-A). For both bacterial species goethite, hematite and calcite showed a lower number of CFUs compared to mica, quartz, and the control (no mineral). These differences have been analyzed statistically using Student t-tests (Fig S4-B) Two of the minerals used in the experiments showed a different effect on the two species. Kaolinite had higher cell viability values for *A. baylyi*, comparable to those found in mica, quartz, and the controls, whereas it had lower cell viability values for *B. subtilis*. However, the viability of *B. subtilis* in kaolinite and *A. baylyi* in calcite is significantly different from both the negatively and positively charged minerals, as it has values in between these two groups (Fig S4). In the case of calcite and *A.baylyi* we have a p-value of 0.0525 when compared to goethite (Supplementary), which is close to the significance level. Further the statistically heat maps cluster the minerals with a similar effect on cell viability. In all cases the clustering follows overall mineral surface charge. For *B. subtilis*, mica and quarts behave similar, and calcite, goethite and hematite have a similar effect on viability. For *A. baylyi*, kaolinite has a similar effect on cell viability the negatively charged minerals and hematite has a distinct effect on viability compared to goethite and quartz which have a similar effect on viability.

**AFM Data**

We imaged the surfaces of the mineral pucks with AFM (Fig S5). The imaging was done in tapping mode in air using a Tap-300 probe with an aluminum reflex coating, a resonant frequency of 300 kH and a force constant of 40 N/m. We calculated the surface roughness and derived the surface area pr unit area (Table S5).

|  | Height image | Phase image | Height image | Phase image |
| --- | --- | --- | --- | --- |
| Calcite | A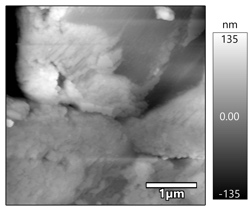 | B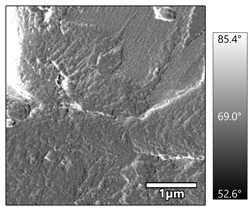 | C 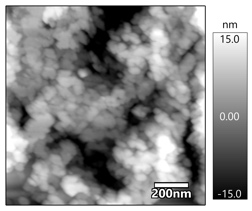 | D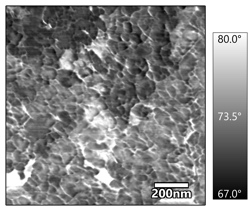 |
| Goethite | 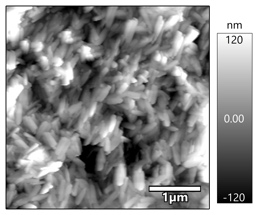 A | 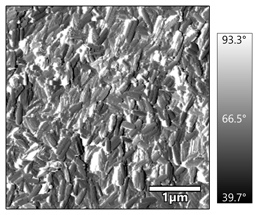 B | 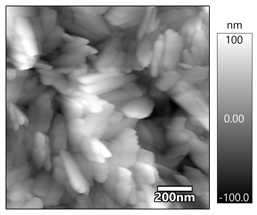 C | 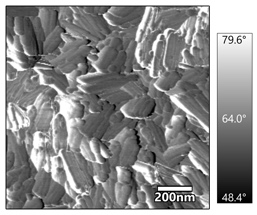 D |
| Hematite | A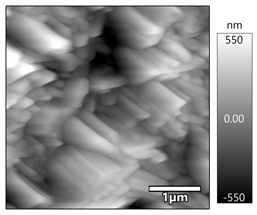 | B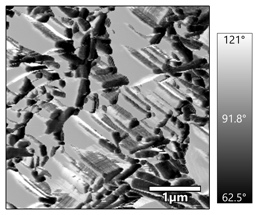 | 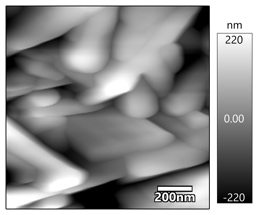 C | 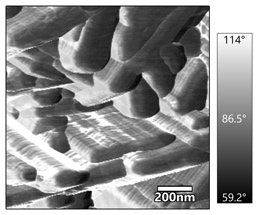 D |
| Kaolinite | 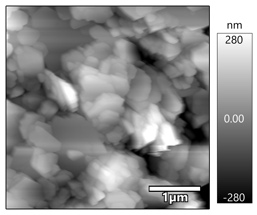 A | 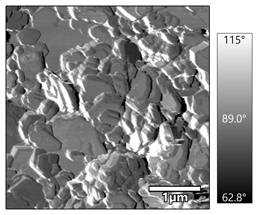 B | 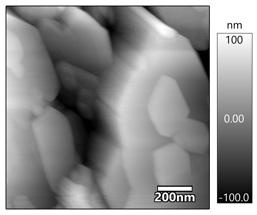 C | 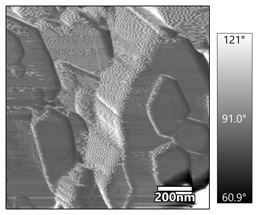 D |
| Mica | A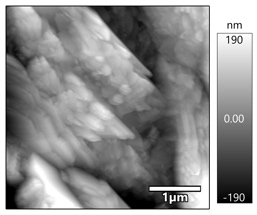 | B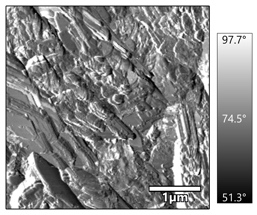 | C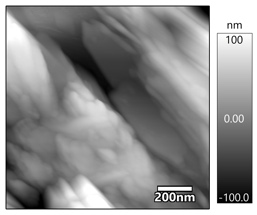 | D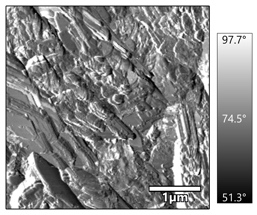 |
| Quartz | A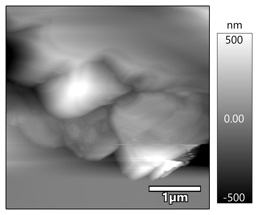 | B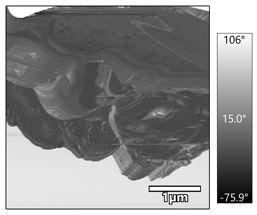 | C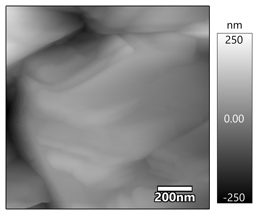 | D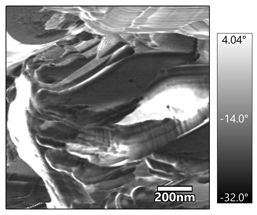 |

**Fig S5.**  Height and phase AFM images of the mineral substrates. Each row corresponds to one mineral, and each column, to either height or phase images which is indicated on top of the figure. A and B show 4 µm resolution images; C and D, 1.20 um resolution images.

**Table S5.** Roughness (nm), surface area (µm^2^) and surface area to area ratio were calculated for all the minerals in two resolutions 4x4 µm and 1.20x1.20 µm.

|  |  | **Roughness (Standard Deviation) (nm)** | **Surface area (µm^2^)** | **Surface area to area ratio** |  | **Roughness (Standard Deviation) (nm)** | **Surface area (µm^2^)** | **Surface area to area ratio** |
| --- | --- | --- | --- | --- | --- | --- | --- | --- |
|  | Img | resolution: 4x4 µm | | | Img | resolution: 1.20x1.20 µm | | |
| **Calcite** | 0009 | 141.864 | 19.64 | 1.228 | 0011 | 27.398 | 1.675 | 1.163 |
|  | 0012 | 68.324 | 17.98 | 1.124 | 0014 | 17.725 | 1.658 | 1.151 |
|  | 0092 | 68.863 | 17.01 | 1.063 | 0095 | 42.405 | 1.668 | 1.158 |
|  | 0096 | 48.947 | 16.88 | 1.055 | 0097 | 7.612 | 1.496 | 1.039 |
|  | 0100 | 85.401 | 20.6 | 1.288 | 0102 | 32.789 | 2.074 | 1.440 |
| Average |  | **82.680** | **18.422** | **1.151** |  | **25.586** | **1.714** | **1.190** |
| **SD*** |  | **31.765** | **1.469** | **0.092** |  | **12.024** | **0.192** | **0.133** |
| **Goethite** | 0004 | 58.845 | 21.66 | 1.354 | 0002 | 37.323 | 2.221 | 1.542 |
|  | 0005 | 53.231 | 20.99 | 1.312 | 0007 | 38.076 | 2.135 | 1.483 |
|  | 0011 | 72.803 | 22.1 | 1.381 | 0016 | 37.601 | 1.944 | 1.350 |
|  | 0015 | 57.015 | 21.26 | 1.329 |  |  |  |  |
| Average |  | **60.474** | **21.503** | **1.344** |  | **37.667** | **2.100** | **1.458** |
| SD |  | **7.401** | **0.419** | **0.026** |  | **0.311** | **0.116** | **0.080** |
| **Hematite** | 0017 | 204.104 | 30.46 | 1.904 | 0019 | 88.245 | 2.485 | 1.726 |
|  | 0018 | 217.205 | 30.39 | 1.899 | 0022 | 126.917 | 2.686 | 1.865 |
|  | 0021 | 276.05 | 35.18 | 2.199 | 0023 | 113.541 | 3.478 | 2.415 |
|  | 0024 | 306.417 | 37.06 | 2.316 |  |  |  |  |
| Average |  | **250.944** | **33.273** | **2.080** |  | **109.568** | **2.883** | **2.002** |
| SD |  | **41.952** | **2.924** | **0.183** |  | **16.036** | **0.429** | **0.298** |
| **Kaolinite** | 0028 | 240.944 | 26.61 | 1.663 |  |  |  |  |
|  | 0029 | 161.418 | 25.6 | 1.600 | 0031 | 40.6 | 1.824 | 1.267 |
|  | 0032 | 132.36 | * | * | 0067 | 19.433 | 1.566 | 1.088 |
|  | 0039 | 99.22 | 21.21 | 1.326 | 0071 | 79.678 | 2.256 | 1.567 |
|  | 0042 | 107.273 | 23.02 | 1.439 |  |  |  |  |
|  | 0066 | 115.714 | 18.48 | 1.155 |  |  |  |  |
| Average |  | **142.822** | **22.984** | **1.437** |  | **46.570** | **1.882** | **1.307** |
| SD |  | **48.275** | **2.947** | **0.184** |  | **24.955** | **0.285** | **0.198** |
| **Mica** | 0024 | 31.083 | 16.7 | 1.044 | 0108 | 8.885 | 1.481 | 1.028 |
|  | 0025 | 49.843 | 18.59 | 1.162 | 0111 | 60.431 | 1.848 | 1.283 |
|  | 0110 | 43.343 | 19.01 | 1.188 | 0114 | 40.18 | 1.79 | 1.243 |
|  | 0112 | 82.916 | 19.27 | 1.204 | 0115 | 54.395 | 1.846 | 1.282 |
|  | 0113 | 101.128 | 19.56 | 1.223 |  |  |  |  |
| Average |  | **61.663** | **18.626** | **1.164** |  | **40.973** | **1.741** | **1.209** |
| SD |  | **26.152** | **1.014** | **0.063** |  | **19.931** | **0.152** | **0.106** |
| **Quartz** | 0000 | 81.318 | 1.337** | 1.181** | 0001 | 42.627 | 1.768 | 1.228 |
|  | 0003 | 159.834 | 11.5** | 1.276** | 0004 | 60.732 | 1.885 | 1.309 |
|  | 0007 | 117.651 | 6.282** | 1.255** | 0008 | 50.499 | 1.954 | 1.357 |
| Average |  | **119.601** | - | **1.238** |  | **51.286** | **1.869** | **1.298** |
| SD |  | **32.084** | - | **0.041** |  | **7.412** | **0.077** | **0.053** |

* Standard deviation (SD)

** On these images, the mineral substrates did not cover completely the scan area. Hence, the surface area values and the surface area to area ratios of the images 0000, 0003 and 0007 were calculated on a smaller frame: 1.132, 9.01 and 5.004 µm^2^ respectively. The roughness values represented as standard deviation (Rq) were calculated using the software Igor Pro 6.38B01. The surface area was calculated using the software Gwyddion.

**HGT**

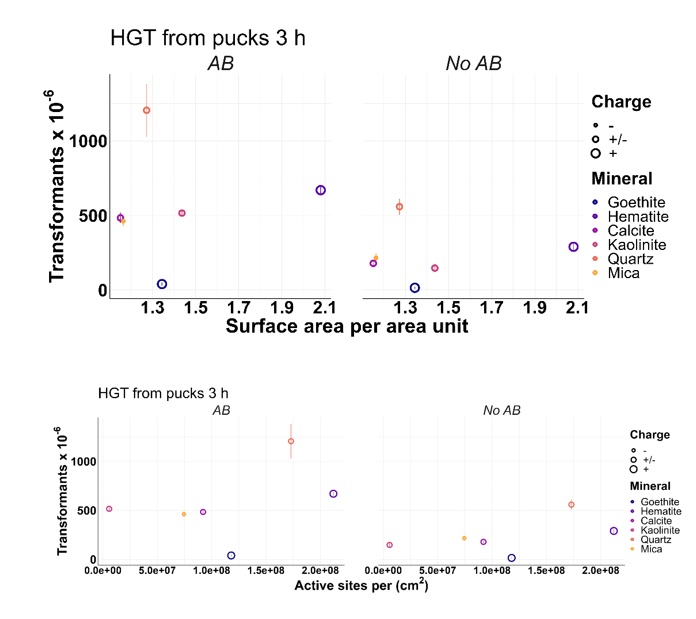

**Figure S6.** **HGT from pucks after 3 hr.** Up) transformation frequencies vs surface area with and without antibiotic, Down) transformation frequencies vs active sites pr area with and without antibiotic. Both show minerals and their respective charges.
